# A unifying mechanism for the biogenesis of prokaryotic membrane proteins co-operatively integrated by the Sec and Tat pathways

**DOI:** 10.1101/111781

**Authors:** Fiona J. Tooke, Marion Babot, Govind Chandra, Grant Buchanan, Tracy Palmer

## Abstract

The vast majority of polytopic membrane proteins are inserted into the cytoplasmic membrane of prokaryotes by the general secretory (Sec) pathway. However, a subset of monotopic proteins that contain non-covalently-bound redox cofactors depend on the twin-arginine translocase (Tat) machinery for membrane integration. Recently actinobacterial Rieske iron-sulfur cluster-containing proteins were identified as an unusual class of membrane proteins that require both the Sec and Tat pathways for the insertion of their three transmembrane domains (TMDs). The Sec pathway inserts the first two TMDs of these proteins co-translationally, but releases the polypeptide prior to the integration of TMD3 to allow folding of the cofactor-containing domain and its translocation by Tat. Here we have investigated features of the *Streptomyces coelicolor* Rieske polypeptide that modulate its interaction with the Sec and Tat machineries. Mutagenesis of a highly conserved loop region between Sec-dependent TMD2 and Tat-dependent TMD3 shows that it plays no significant role in coordinating the activities of the two translocases, but that a minimum loop length of approximately eight amino acids is required for the Tat machinery to recognise TMD3. Instead we show that a combination of relatively low hydrophobicity of TMD3, coupled with the presence of C-terminal positively-charged amino acids, results in abortive insertion of TMD3 by the Sec pathway and its release at the cytoplasmic side of the membrane. Bioinformatic analysis identified two further families of polytopic membrane proteins that share features of dual Sec-Tat-targeted membrane proteins. A predicted heme-molybdenum cofactor-containing protein with five TMDs, and a polyferredoxin also with five predicted TMDs, are encoded across bacterial and archaeal genomes. We demonstrate that membrane insertion of representatives of each of these newly-identified protein families is dependent on more than one protein translocase, with the Tat machinery recognising TMD5. Importantly, the combination of low hydrophobicity of the final TMD and the presence of multiple C-terminal positive charges that serve as critical Sec-release features for the actinobacterial Rieske protein also dictate Sec release in these further protein families. Therefore we conclude that a simple unifying mechanism governs the assembly of dual targeted membrane proteins.

## Introduction

Prokaryotic cytoplasmic membrane proteins represent 20-30% of the proteome (1, 2) and they fulfil a wide variety of critical functions in the cell including respiration, photosynthesis, and ion transport, allowing this membrane to act as a tightly controlled barrier between the cytoplasm and the extracellular environment. Cytoplasmic integral membrane proteins adopt α-helical topologies, and in bacteria are inserted via the action of at least one of three protein translocation machineries - the Sec machinery, the YidC insertase and the Tat pathway (see (3) for a recent review).

The SecYEG translocon is the major route by which multi-spanning membrane proteins are integrated into the membrane. The insertion of transmembrane domains of polytopic proteins occurs co-translationally following targeting of the translating ribosome to the Sec machinery through the action of signal recognition particle (SRP) (4). YidC is positioned close to the lateral gate of SecY and interacts with nascent transmembrane domains to facilitate their integration into the membrane (5-7). YidC can also act independently of the Sec system to integrate small (usually mono- or bitopic) membrane proteins directly into the bilayer (8, 9). The final topology adopted by a polytopic membrane protein depends upon a number of intrinsic and extrinsic factors including the hydrophobicity of membrane-spanning regions, the number and location of positively-charged amino acids and the composition of the lipid bilayer (10-12).

The Tat system is a post-translational protein transport pathway that operates independently of the Sec and YidC machineries to transport folded proteins across the cytoplasmic membrane (reviewed in 13, 14). Proteins are targeted to the Tat machinery by N-terminal signal sequences containing a highly conserved pair of arginine residues that are usually critical for efficient recognition of substrates (15). A subset of Tat substrate proteins contain non-covalently bound prosthetic groups such as metal-sulphur clusters or nucleotide-based cofactors, many of which play important roles in respiratory and photosynthetic metabolism (16). Some Tat substrates are also integral membrane proteins. In bacteria Tat-dependent integral membrane proteins generally fall into two classes – those that are N-terminally anchored in the bilayer by a non-cleaved signal sequence, such as the Rieske iron-sulfur proteins for example of *Paracoccus* or *Legionella* (17, 18) or the TtrA subunit of *Salmonella* tetrathionate reductase (19) and those that have a single transmembrane helix at their C-termini such as the small subunits of hydrogenases and formate dehydrogenases (20, 21). Recent studies have indicated that the Rieske proteins of actinobacteria are highly unusual Tat substrates (22, 23). Rieske proteins are essential membrane-bound components of cytochrome *bc_1_* and *b_6_f* complexes that coordinate an iron-sulfur (FeS) cluster involved in the electron transfer from quinones to cytochromes *c_1_/f* (for reviews see (24, 25)). The actinobacterial proteins have three transmembrane domains (TMDs) preceding the Rieske FeS domain, unlike most other Rieske proteins which contain only one TMD. Inspection of actinobacterial Rieske sequences indicates the presence of a predicted twin-arginine motif between TMDs 2 and 3, suggesting the possibility that the concerted action of more than one translocase may be required for correct assembly. Indeed it was shown that the first two TMDs of the *Streptomyces coelicolor* Rieske protein, Sco2149, are inserted by the Sec machinery, probably in a co-translational manner, whereas the insertion of TMD3 is dependent on the Tat pathway (22), providing the first example of these two machineries operating together to assemble a single protein.

These findings raise a number of pertinent questions about the mechanisms by which these translocases are co-ordinated to ensure that the Sec system does not integrate TMD3 but releases the polypeptide to allow folding of the globular domain, and the subsequent recognition of a membrane-tethered substrate by the Tat pathway. It also raises the question whether actinobacterial Rieske proteins represent an oddity of nature, or whether there are further examples of dual Sec/Tat-targeted membrane proteins to be discovered. Here we have addressed both of these major aspects and show that in addition to Rieske there are at least two further conserved families of dual targeted membrane proteins across bacteria and archaea that each have 5 TMDs. A detailed dissection of the features of the transmembrane regions of *S. coelicolor* Rieske reveals that the relatively low hydrophobicity of TMD3 coupled with the location of positively charged amino acid residues orchestrate the release of the polypeptide by the Sec pathway. Importantly, we demonstrate that these features are also present across all identified families of these dual-targeted membrane proteins indicating that there is unifying mechanism for their biogenesis.

## Results

### Fusion proteins for the analysis of Sco2149 membrane assembly

Previous work has shown that the *S. coelicolor* Rieske protein, Sco2149, has three transmembrane domains that require the combined action of two distinct protein translocases, Sec and Tat, for complete assembly into the membrane (22, 23). However the mechanism by which these two translocases are coordinated is unknown, although TMD and globular domain swapping experiments indicated that the information required to coordinate this process does not reside within the first two TMDs or the cofactor binding domain (22).

To assess the mechanism of TMD insertion we used constructs where the cofactor-containing FeS domain was genetically removed from Sco2149 and replaced with the mature region of two different reporter proteins – that of the *E. coli* Tat substrate AmiA (26) to report on interaction of Sco2149 with the Tat pathway, or of the Sec substrate β-lactamase (Bla, which is compatible for export with either the Sec or Tat pathways depending on the nature of the targeting sequence (27)) (Fig 1A, Fig S1). These constructs were produced from the medium copy number vector pSU-PROM (which specifies kanamycin resistance (28)) under control of the constitutive *tatA* promoter (29).

**Figure 1:**
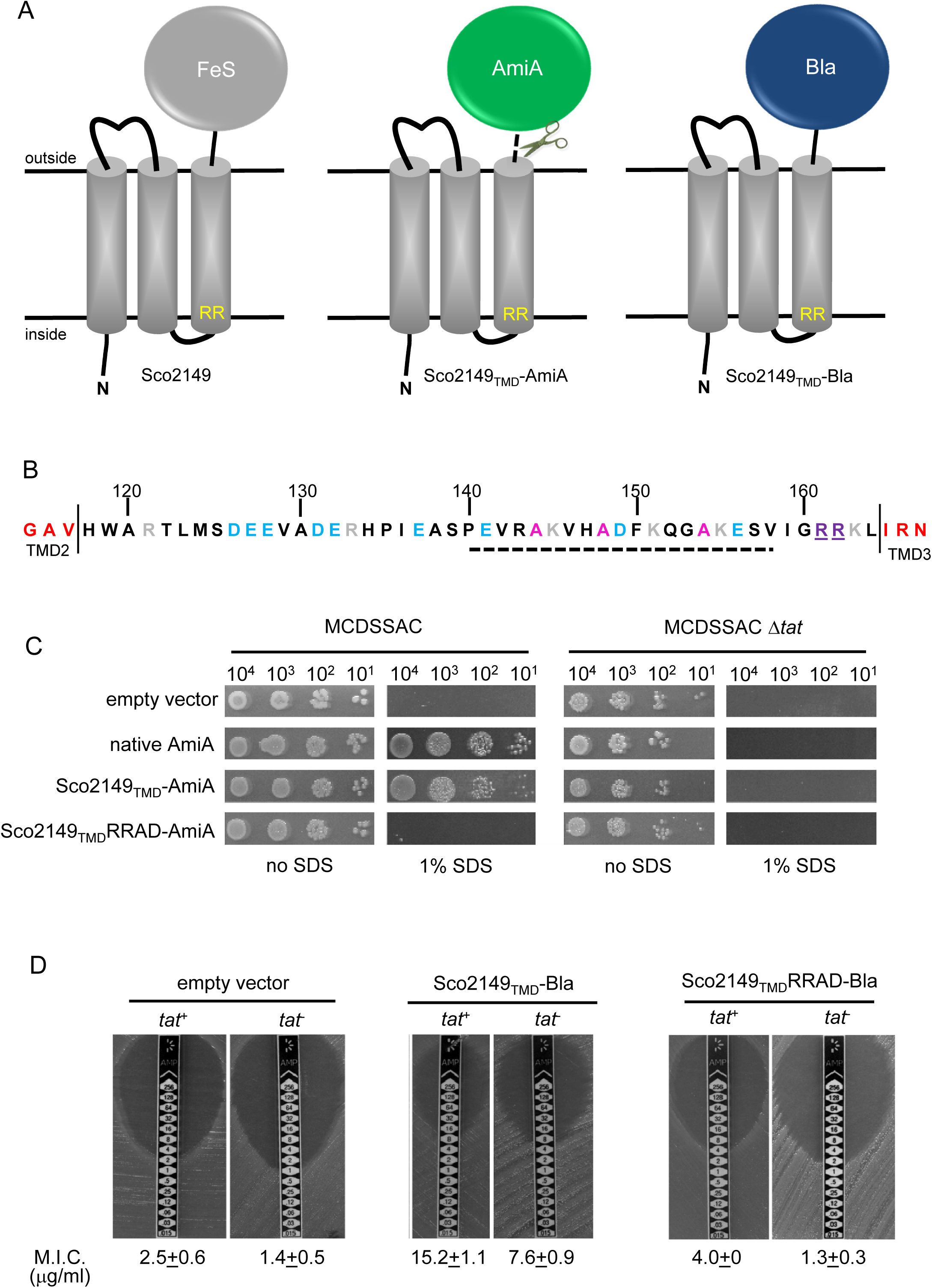
Sco2149_TMD_-reporter fusions to follow membrane insertion. (A) Cartoon representations of the *S. coelicolor* Rieske protein, Sco2149, and the Sco2149_TMD_-AmiA and Sco2149_TMD_-Bla fusions. A signal peptidase I cleavage site (indicated by scissors) was introduced into the between the end of TMD3 and the AmiA sequence to allow release of AmiA from the membrane (22). The position of the twin-arginine motif is indicated by RR. (B) Sequence of the Sco2149 cytoplasmic loop region between TMDs 2 and 3. Amino acids predicted to be part of TMDs 2 and 3 are shown in red. The twin arginines of the Tat recognition motif are given in purple underline. Predicted α-helical secondary structure is shown with a dotted line, and alanine residues within this region that were mutated to proline are shown in pink. Negatively charged amino acids in the loop region are shown in blue, positively charged ones in grey. (C) *E. coli* strain MCDSSAC (which carries chromosomal deletions in the signal peptide coding regions of *amiA* and *amiC)* or an isogenic *tatABC* mutant containing either pSU-PROM (empty vector), or pSU-PROM producing native AmiA, Sco2149_TMD_-AmiA or a variant where the twin-arginines were substituted to AD (Sco2149_TMD_RRAD-AmiA), were spotted after serial dilution on LB medium in absence or presence of 1% SDS. The plates were incubated for 20 hours at 37°C. (D) Representative images of M.I.C.Evaluator™ strip tests of strains MC4100 (*tat^+^*) and DADE (*tat^-^*) harbouring pSU-PROM (empty vector), pSU-PROM Sco2149_TMD_-Bla or pSU-PROM Sco2149_TMD_RRAD-Bla. The mean M.I.C ± s.d. for strains harbouring these constructs is given at the bottom of each test strip (where *n*=4 biological replicates for each strain harbouring the empty vector, *n=5* biological replicates for each strain harbouring pSU-PROM Sco2149_TM_D-Bla and *n*=3 biological replicates for each strain harbouring pSU-PROM Sco2149_TMD_RRAD-Bla).

AmiA and its homologue AmiC are periplasmic Tat substrates that remodel the peptidoglycan, and in their absence *E. coli* is sensitive to growth in the presence of SDS (26, 30) (Fig 1C; top panel). As expected, when either plasmid-encoded native AmiA or the Sco2149_TMD_-AmiA fusion was produced in the *tat*^+^ strain lacking chromosomally encoded periplasmic AmiA and AmiC (MCDSSAC), growth on SDS was restored (Fig 1C, middle two panels). The export of AmiA from both of these constructs was absolutely dependent on the Tat pathway as no growth on SDS was conferred in the *tat*^-^ strain (MCDSSAC Δ*tat*). Previously it has been reported that a twin lysine substitution of the twin arginine motif of Sco2149 was sufficient to prevent Tat-dependent export of AmiA when produced at lower levels from the pSU18 plasmid (22). However, when expressed from the pSU-PROM vector, a low level of export by the Tat pathway could still be observed for the Sco2149-AmiA construct harbouring this substitution Fig S2). It has been noted previously that Tat-dependent export of some very sensitive plasmid-borne reporter proteins can be detected following twin lysine substitution of the twin arginines (31, 32), indicating that twin lysines can still trigger Tat-dependent export but with a greatly reduced efficiency. However, less conservative substitutions of the twin arginine motif to twin alanine or to alanine-aspartate were not permissive for Tat transport (Fig 1C; Fig S2).

The membrane insertion of Sco2149 was further investigated using the Bla fusion construct. When exported to the periplasmic side of the membrane Bla confers resistance to ampicillin, which can be assessed in a quantitative manner using M.I.C.Evaluator™ test strips. Fig 1D shows that the basal M.I.C. for ampicillin was evaluated at 2.5 and 1.4 μg/ml, respectively, for the *tat*^+^(MC4100) and *tat*^-^ (DADE) strains harbouring the empty vector. We assign these slight differences in M.I.C. to the partially compromised cell wall in *tat* mutant strains (26,30). The *tat*^+^strain producing the Sco2149TMD-Bla fusion protein was able to grow up to a concentration of approximately 15 μg/ml ampicillin, indicating that there was export of Bla in this strain. However, some of that export was clearly by the Sec pathway since the *tat* strain producing Sco2149_TMD_-Bla had an M.I.C. for ampicillin of 7.6 μg/ml, significantly above basal level. It has been reported that the introduction of negative charges into the n-region of a Sec signal peptide blocks Sec-dependent translocation (33), and therefore substituting the twin arginines to alanine-aspartate would be expected to prevent translocation through both the Sec and Tat pathways. As shown in Fig 1D these substitutions reduced the MIC for ampicillin to 4.0 and 1.3 μg/ml, respectively, for *tat*^+^ and *tat*^-^ strain, very close to basal level. Taken together these results indicate that there is some compatibility of TMD3 of the *S. coelicolor* Rieske protein with the Sec pathway, which was not seen previously using a more qualitative assay (22).

### The cytoplasmic loop region of Sco2149 does not modulate interaction of TMD3 with the Sec pathway

The finding that there is some Sec-dependent translocation of the Bla portion of the Sco2149TMD-Bla fusion in a strain lacking the Tat pathway provides a useful tool to study features of the protein that influence interaction with the Sec machinery. We therefore undertook a programme of mutagenesis on the Sco2149_TMD_-Bla construct, focusing firstly on the cytoplasmic loop region between TMD2 and TMD3 as this has a number of highly conserved features across actinobacterial Rieske proteins (Fig 1B; Fig S3). In particular the loop has a highly conserved length (43 amino acids between the predicted end of TM2 and the twin arginine motif), a region of predicted α-helical structure, and a number of positions where positively or negatively charged residues are conserved, including an almost invariant glutamic acid (E127 in Sco2149) and arginine-histidine pairing (R133, H134 in Sco2149).

Initial site-directed replacement of amino acids in the loop region were undertaken and the level of resistance to ampicillin mediated by the variant Sco2149_TMD_-Bla fusion protein in a *tat*^-^ background was scored. As shown in Table 1, apart from the introduction of an alanine-aspartate pair to replace the twin arginines, none of the substitutions we introduced, including replacement of the highly conserved E127 or R133/H134 residues or introduction of proline residues into the predicted α-helical region, had any substantive effect on the interaction of Sco2149 with the Sec pathway. We therefore made further substitutions, for example progressively deleting clusters of negatively charged amino acids or changing them to positively charged lysines. None of these deletions or substitutions had any detectable effect on Sec translocation of the Bla fusion, even when all of the acidic residues were substituted for lysine. Moreover, insertion of three additional negative charges into the loop was also without detectable effect.

**Table 1.** Effect of amino acid substitutions, small deletions and insertions in the Sco2149 cytoplasmic loop region on the ability of Sco2149_TMD_-AmiA and Sco2149_TMD_-Bla to support growth on SDS or ampicillin, respectively. Note that growth on ampicillin was scored using the *tat*^-^ strain DADE and therefore assesses Sec translocation only. Y indicates growth on 1% SDS, N indicates no growth, nd – not determined. Mean M.I.C for growth on ampicillin is given in μg/mI ± one standard deviation, *n* = at least 3. *Insertion of 3 additional amino acids, DEE between E128 and V129.

| Variant | Growth on 1% SDS<br>(Tat translocation) | mean M.I.C. for ampicillin<br>(Sec translocation) |
| --- | --- | --- |
| wild type | Y | 7.6 ± 0.9 |
| R161K R162K | Y | 3.8 ± 0.5 |
| R161A R162D | N | 1.3 ± 0.3 |
| R161A R162A | N | 5.5 ± 1.0 |
| R161K R162Q | N | 3.3 ± 0.6 |
| ΔR161 ΔR162 | N | 9.3 ± 2.3 |
| R133H H134R | Y | 7.3 ± 1.2 |
| R133K H134K | Y | 7.0 ± 2.0 |
| M124L | Y | 6.0 ± 2.0 |
| M124A | Y | 6.0 ± 2.0 |
| S125L | Y | 6.7 ± 2.3 |
| S125A | Y | 8.0 ± 0.0 |
| D126L | Y | 7.3 ± 1.2 |
| D126A | Y | 7.3 ± 1.2 |
| E127L | Y | 6.0 ± 2.0 |
| E127A | Y | 8.0 ± 2.8 |
| A144P | Y | 6.7 ± 1.2 |
| A148P | Y | 8.0 ± 2.8 |
| A154P | Y | 8.0 ± 0.0 |
| Δ126-8 | nd | 8.0 ± 0.0 |
| Δ126-127 | nd | 7.5 ± 1.0 |
| Δ127-128 | nd | 8.0 ± 2.8 |
| Δ131-2 | nd | 7.3 ± 1.2 |
| Δ137Δ141 | nd | 4.0 ± 0.0 |
| Δ131-2Δ141 | nd | 6.0 ± 2.0 |
| Δ149Δ156 | nd | 8.0 ± 0.0 |
| Δ126-8 Δ137Δ141 | nd | 8.0 ± 0.0 |
| Δ126-8Δ131-2 | nd | 8.0 ± 2.4 |
| Δ126-8Δ131-2 Δ137Δ141 | nd | 10.4 ± 2.2 |
| Δ126-8Δ131-2 Δ137 Δ141 Δ149 Δ156 | nd | 10.0 ± 2.8 |
| Ins D129 E130 E131* | nd | 6.7 ± 1.2 |
| D131A E132A | nd | 6.0 ± 2.3 |
| E137A E141A | nd | 6.7 ± 2.3 |
| D126A E127A E128A | nd | 7.2 ± 1.1 |
| D131K E132K | nd | 7.3 ± 1.2 |
| E137K E141K | nd | 6.5 ± 1.9 |
| D126K E127K E128K | nd | 9.0 ± 2.0 |
| D126K E127K E128K E137K E141K | nd | 8.0 ± 0.0 |
| D126K E127K E128K D131K E132K<br>E137K E141K | nd | 10.0 ± 2.2 |

We similarly assessed translocation by Sec for a series of sliding truncations of 5, 10, 15, 20, 25, 30 and 35 residues within the loop region (summarised in Table 2). Again most of the truncations had little effect on translocation of Sco2149_TMD_-Bla by the Sec pathway, and even truncations of 30 residues or more gave mean M.I.C.s for ampicillin similar to that seen for the non-mutated construct. These findings indicate that many of the conserved features noted in this loop region, for example the overall length, presence of a predicted α-helical region and clusters of negatively charged amino acids do not modulate interaction of Sco2149 with the Sec pathway.

**Table 2.**
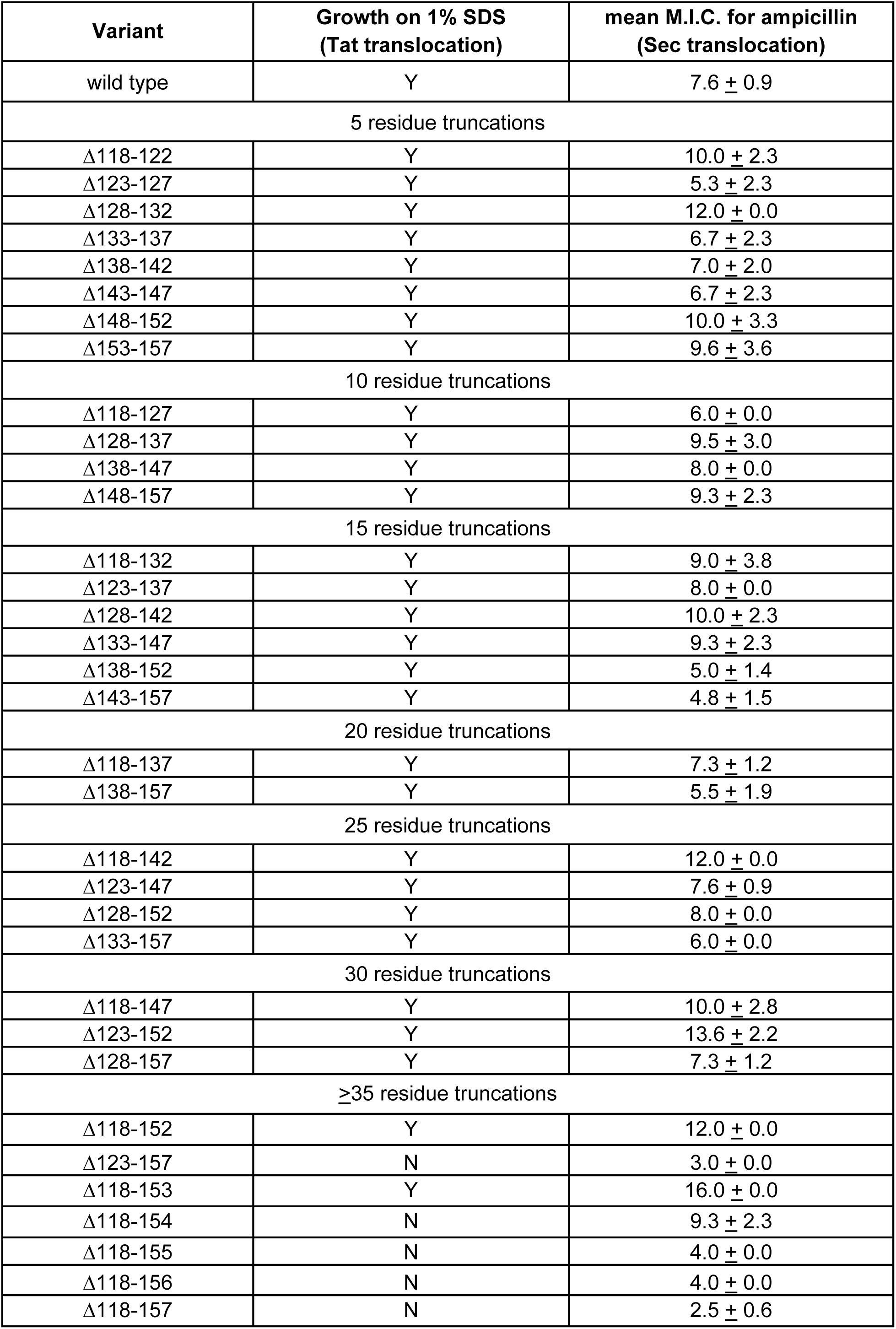
Effect of amino acid truncation in the Sco2149 cytoplasmic loop region on the ability of Sco2149_TMD_-AmiA and Sco2149_TMD_-Bla to support growth on SDS or ampicillin, respectively. Note that growth on ampicillin was scored using the *tat* strain DADE and therefore assesses Sec translocation only. Y indicates growth on 1% SDS, N indicates no growth. Mean M.I.C for growth on ampicillin is given in μg/ml ± one standard deviation, *n* = at least 3.

| Variant | Growth on 1% SDS<br>(Tat translocation) | mean M.I.C. for ampicillin<br>(Sec translocation) |
| --- | --- | --- |
| wild type | Y | 7.6 ± 0.9 |
| 5 residue truncations |  |  |
| Δ118-122 | Y | 10.0 ± 2.3 |
| Δ123-127 | Y | 5.3 ± 2.3 |
| Δ128-132 | Y | 12.0 ± 0.0 |
| Δ133-137 | Y | 6.7 ± 2.3 |
| Δ138-142 | Y | 7.0 ± 2.0 |
| Δ143-147 | Y | 6.7 ± 2.3 |
| Δ148-152 | Y | 10.0 ± 3.3 |
| Δ153-157 | Y | 9.6 ± 3.6 |
| 10 residue truncations |  |  |
| Δ118-127 | Y | 6.0 ± 0.0 |
| Δ128-137 | Y | 9.5 ± 3.0 |
| Δ138-147 | Y | 8.0 ± 0.0 |
| Δ148-157 | Y | 9.3 ± 2.3 |
| 15 residue truncations |  |  |
| Δ118-132 | Y | 9.0 ± 3.8 |
| Δ123-137 | Y | 8.0 ± 0.0 |
| Δ128-142 | Y | 10.0 ± 2.3 |
| Δ133-147 | Y | 9.3 ± 2.3 |
| Δ138-152 | Y | 5.0 ± 1.4 |
| Δ143-157 | Y | 4.8 ± 1.5 |
| 20 residue truncations |  |  |
| Δ118-137 | Y | 7.3 ± 1.2 |
| Δ138-157 | Y | 5.5 ± 1.9 |
| 25 residue truncations |  |  |
| Δ118-142 | Y | 12.0 ± 0.0 |
| Δ123-147 | Y | 7.6 ± 0.9 |
| Δ128-152 | Y | 8.0 ± 0.0 |
| Δ133-157 | Y | 6.0 ± 0.0 |
| 30 residue truncations |  |  |
| Δ118-147 | Y | 10.0 ± 2.8 |
| Δ123-152 | Y | 13.6 ± 2.2 |
| Δ128-157 | Y | 7.3 ± 1.2 |
| ≥35 residue truncations |  |  |
| Δ118-152 | Y | 12.0 ± 0.0 |
| Δ123-157 | N | 3.0 ± 0.0 |
| Δ118-153 | Y | 16.0 ± 0.0 |
| Δ118-154 | N | 9.3 ± 2.3 |
| Δ118-155 | N | 4.0 ± 0.0 |
| Δ118-156 | N | 4.0 ± 0.0 |
| Δ118-157 | N | 2.5 ± 0.6 |

We did note, however, that one of the 35 residue truncations, Δ123-157, significantly reduced integration of TMD3 by the Sec pathway (Fig 2A,B), whereas the other 35 residue truncation, Δ118-152, showed a slight increase in Sec translocation (c.f. M.I.C of 7.6 μg/ml ampicillin for the non-mutated construct vs 12 μg/ml for the Δ118-152 truncation). This suggested that there may be some feature of the loop region between residues 153 and 157 influencing interaction with the Sec pathway. To explore this further we made a series of additional one amino acid truncations to give Δ118-153, Δ118-154, Δ118-155 and Δ118-156 and Δ118-157 constructs. Fig 2B indicates that as soon as the truncation extended to amino acid 155, Sec translocation was substantially reduced (but protein production and/or stability was not, Fig 2C). Inspection of the sequence indicates that amino acid 155 is a lysine. Positively charged amino acids are important topology determinants in membrane proteins, and are enriched in the cytoplasmic regions of membrane proteins, the so-called ‘positive inside rule’ due to the energetic cost of translocating them across the membrane against the protonmotive force (34, 35). To test whether the loss of this basic residue was the reason for the very low level of periplasmic Bla activity, we introduced a positive charge further along the loop (V158K) into the full length Sco2149_TMD_-Bla and the Δ118-155, Δ118-156 and Δ118-157 truncations. Fig 2D shows that the introduction of the V158K into the Δ118-155, Δ118-156 and Δ118-157 truncations restored the M.I.C. to a similar level seen for the full length Sco2149_TMD_-Bla, establishing that positive charged residues in this loop region influence interaction of Sco2149 with Sec.

**Figure 2:**
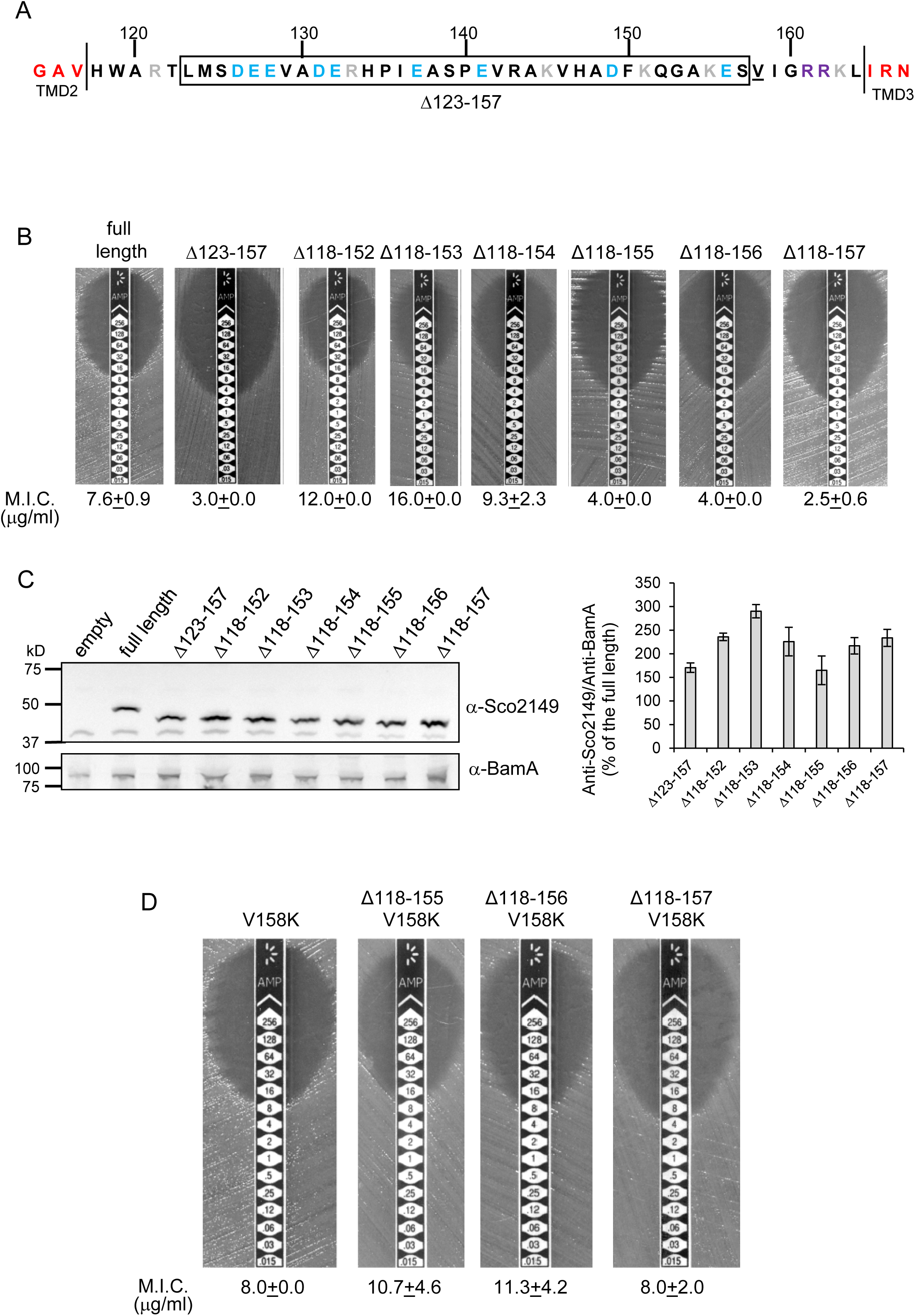
Interaction of Sco2149_TMD_-Bla with the Sec pathway. (A) The Sco2149 cytoplasmic loop region between TMDs 2 and 3. Color-coding is as described in Fig 1. The extent of the 123-157 deletion is shown boxed and V158 that was substituted to K in this study is underlined. (B) and (D) Representative images of M.I.C.Evaluator™ strip tests of strain DADE (*tat*) harbouring pSU-PROM producing the indicated variants of Sco2149_TMD_-Bla. The mean M.I.C ± s.d. is given at the bottom of each test strip (where *n*=3 biological replicates for each strain). (C) Membrane extracts prepared from the same strains used in (B) along with DADE harboring the empty plasmid vector as a negative control, were separated by SDS-PAGE (12% acrylamide), transferred to nitrocellulose membrane and probed with anti-Sco2149 or anti-BamA (an unrelated membrane protein was used as a loading control). To the right, the Sco2149-associated signal was quantified and normalised against the BamA signal for each sample. The quantification results were expressed as percentage of the normalised signal obtained for the full length fusion (which was set at 100%). The results represent mean ± s.e.m. of three biological replicates.

### A minimum cytoplasmic loop length is necessary for Tat recognition of Sco2149 TMD3

Since none of the conserved features in the Sco2149 cytoplasmic loop were required for modulating interaction with the Sec pathway, we next addressed whether they were required for recognition by the Tat system. A subset of the amino acid substitutions and each of the sliding truncations was introduced into the Sco2149_TMD_-AmiA fusion protein and expressed in a *tat*^+^ strain to allow Tat-dependence to be scored by testing for growth in the presence of SDS (Tables 1, 2). Table 1 shows that, apart from substitutions at the twin arginine motif, none of the other variants affected Tat-dependent export of AmiA, including the introduction of prolines within the predicted α-helical structure, or substitution of the highly conserved E127 or R133/H134. These results suggest that none of these features are required for recognition of the loop region by the Tat pathway.

Ordinarily, Tat signal peptides have free N-termini, whereas the Tat signal sequence of Sco2149 is internal and is only recognised by the Tat pathway once the first 2 TMD of the protein have been integrated by Sec. The loop truncation experiments indicated that the Tat system was still able to identify and integrate TMD3 when it was truncated by up to 30 residues. However, one of the 35 residue truncations (Sco2149_TMD_-Δ123-157-AmiA) and the 40 residue truncation (Sco2149_TMD_-Δ118-157-AmiA) supported no growth on SDS-containing media (Table 2), indicating that there is a minimum loop length requirement of approximately eight amino acids between TMD2 and the twin arginine motif is required for Tat recognition of a tethered signal peptide.

Taken together we conclude that, with the exception of the twin arginine motif, none of the conserved features of cytoplasmic loop are strictly necessary for interaction of Sco2149 with the Tat pathway or to mediate release from Sec.

### Specific physical properties of TMD3 drive its release from Sec

Hydrophobicity is the driving force for the insertion of a helix into the membrane (10, 36, 37). Analysis of transmembrane helices from polytopic proteins of known three-dimensional structure shows a general trend that the first and last TMDs are of similar hydrophobicity, and they are notably more hydrophobic than the central helices (38, 39). An analysis of the apparent ΔG for the insertion of the three TMDs of selected actinobacterial Rieske proteins is shown in Table 3. It can be seen that the first and second TMDs have negative predicted ΔG_app_ values and are therefore expected to be inserted as TMDs by the Sec system (40). However, the third and final TMD is predicted to have a positive Gapp (Table 3). This is in contrast to the final TMD of ‘standard’ Sec-dependent proteins and suggests that this helix might be poorly recognised by the Sec machinery.

**Table 3.** Predicted ΔG_app_ values (in kcal mol^-1^) for membrane insertion of each of the three TMDs of the indicated Rieske proteins analysed using the ΔG_app_ prediction server (http://dgpred.cbr.su.se/) and based on hydrophobicity scales generated from (36, 69). This server uses the SCAMPI2/TOPCONS servers (67, 68) to predict the positions of the TMDs and for *S. coelicolor* Rieske predicts TMD1 to span aa 58-80, TMD 2 to span aa 96-117 and TMD3 to span aa 168-187.

| Family/Species | Uniprot ID | Predicted $\Delta G_{app}$ | | | TM3/Bla fusion* |
| --- | --- | --- | --- | --- | --- |
|  |  | TM1 | TM2 | TM3 |  |
| *Mycobacterium tuberculosis* | P9WH23 | -4.252 | -0.715 | 0.248 |  |
| *Corynebacterium glutamicum* | Q79VE8 | -2.873 | -0.761 | 0.564 |  |
| *Gordonia malaquae* | M3VAA9 | -2.569 | -0.214 | 1.291 |  |
| *Corynebacterium diphtheriae* | Q6NGA2 | -2.402 | -1.089 | 0.619 |  |
| *Dietzia cinnamomea* | E6JC04 | -2.997 | -0.391 | 0.409 |  |
| *Salinispora tropica* | A4X9Y7 | -2.315 | -1.205 | 1.162 |  |
| *Streptomyces* sp. | D9VGG2 | -1.291 | -1.203 | 0.967 |  |
| *Verrucospora maris* | F4F1U1 | -2.248 | -1.946 | 1.162 |  |
| *Stackebrandtia nassauensis* | D3Q119 | -1.876 | -1.289 | 0.556 |  |
| *Rhodococcus erythropolis* | C3JJ95 | -2.692 | -0.208 | 0.912 |  |
| *Streptomyces coelicolor* | Q9X807 | -2.374 | -0.117 | 0.614 | 0.714 |
| *S. coelicolor* S179L |  |  |  | -0.389 | -0.205 |
| *S. coelicolor* G180L |  |  |  | -0.037 | 0.044 |
| *S. coelicolor* S179L, G180L |  |  |  | -1.117 | -0.987 |
| *S. coelicolor* P177L, S179L, G180L |  |  |  | -2.510 | -2.409 |
| *S. coelicolor* R185A |  |  |  | 0.517 | 0.620 |

\*Bla is fused to *S. coelicolor* Rieske after amino acid 185 (full sequence of all of the fusion proteins used in this study can be found in Supplementary Information).

To probe this further we investigated the effect of increasing the hydrophobicity of TMD3. Table 3 shows that substitution of a single leucine residue at either serine 179 or glycine 180 reduces the predicted ΔG_app_ value for TMD3 Sec-dependent membrane insertion by at least 0.6 kcal mol^-1^ Accordingly, when these single substitutions were individually introduced into the Sco2149_TMD_-Bla fusion in *tat*^-^ cells, a dramatic increase in M.I.C for ampicillin of up to 25 fold was observed (Fig 3B), almost at the upper limit of detection. Combining these substitutions (S179L, G180L), and including a third substitution (P177L) shifts the predicted ΔG_app_ value closer to that of TMD1 (Table 1). These substitutions also significantly increased the observed M.I.C. over the unsubstituted fusion, but did not appear to have additive effects over the single leucine substitutions. We conclude that the low hydrophobicity of TMD3 is a key driver for the release of Sco2149 from the Sec machinery.

**Figure 3:**
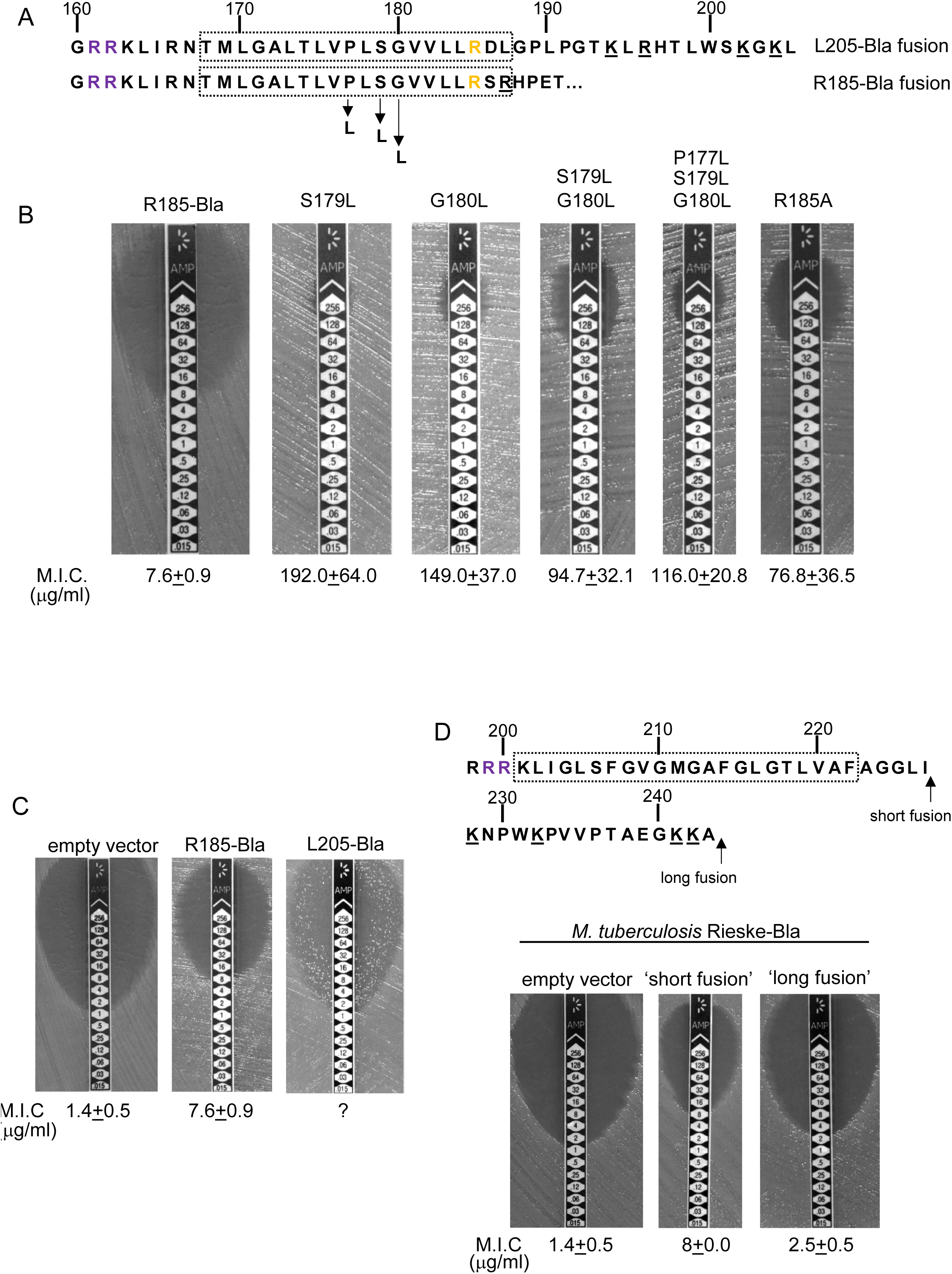
Hydrophobicity of TMD3 and C-terminal positive charges modulate interaction with the Sec pathway. (A) Sco2149 TMD3 and flanking sequences. *Top* shows the native Sco2149 sequence up to amino acid 205 (position of the L205-Bla fusion) and *below* the sequence of the R185 Sco2149-Bla fusion. In each case the predicted position of TMD3 was determined using the SCAMPI2/TOPCONS servers (67, 68) and is shown boxed. The twin arginines are shown in purple and R185 (the position after which Bla was fused in the R185 construct) is shown in yellow. Positively charged amino acids C-terminal to R185 are underlined. (B-D) Representative images of M.I.C.Evaluator™ strip tests of strain DADE (*tat*) harbouring pSU-PROM producing (B) the indicated variants of the R185 Sco2149_TMD_-Bla fusion or (C) the R185 or L205 Sco2149_TMD_-Bla fusions, as indicated, or (D) *M. tuberculosis* QcrA fused to Bla. In (D) the amino acid sequence around TMD3 of *M. tuberculosis* QcrA is shown, with TMD3 boxed. ‘short fusion’ refers to a Bla fusion after I227 and ‘long fusion’ to a Bla fusion after A243. In each panel the mean M.I.C ± s.d. is given at the bottom of each test strip (where *n*=3 biological replicates for each strain).

It has long been known that Tat signal peptides frequently contain one or more positive charges in their c-regions, close to the site of signal peptidase cleavage. These charges are not required for the interaction with the Tat pathway but reduce the efficiency of interaction with Sec and have therefore been described ‘Sec-avoidance’ motifs (41-43). A positive charge is generally also found close to the C-terminal end of TMD3 of actinobacterial Rieske proteins (R185 in the case of Sco2149; Fig 3A, Fig S1). Substitution of R185 for alanine in the Sco2149_TMD_-Bla fusion conferred an 8-fold increase in M.I.C for ampicillin, and therefore R185 also appears to act as a Sec-avoidance motif in this context. Interestingly, closer inspection of actinobacterial Rieske proteins indicates that there are a number of further non-conserved positive charges located within the C-terminal vicinity of TMD3 (Fig 3A underlined residues, Fig S1 orange residues) which are not found in other Rieske proteins that only contain a single TMD (Fig S1B). Since our original Sco2149TMD-Bla fusion (where the Bla sequence is fused immediately after R185) lacks most of these additional charges (Fig 3A), we made an additional Bla fusion where the Sco2149 sequence in the fusion protein was extended to aa205, incorporating an additional four positively charged residues. It can be seen that inclusion of this additional positively charged stretch almost completely abolished transport via Sec, as the clearance zone around the M.I.C. strip was of similar size to that of the negative control (Fig 3C). We did, however, note that for unknown reasons there was a variable level of breakthrough growth within the zone of clearing for strain DADE producing the extended Sco2149_TMD_-Bla fusion. We therefore constructed similar Bla fusions after TMD3 of the *M. tuberculosis* Rieske protein, QcrA. Fig 3D indicates that there is some Sec-dependent export of the Bla fusion when it is fused close to the C-terminal end (‘short fusion’) but that this was almost abolished when the sequence was extended to introduce the positively charged stretch (‘long fusion’). Taken together, we conclude that a combination of low hydrophobicity of TMD3 coupled with the presence of several C-terminal positive charges promotes release of actinobacterial Rieske proteins from the Sec machinery.

### Bioinformatic analysis identifies further families of membrane proteins potentially dependent on both Sec and Tat pathways

We next asked whether actinobacterial Rieske proteins were the only protein family that required both Sec and Tat pathways for their integration. To this end, all proteins from prokaryotic genomes available in Genbank were analysed by both TATFind 1.4 (44) and TMHMM 2.0c (2) programs. For each protein, both outputs were combined to identify the position of twin arginine motif, and the number of transmembrane helices present N-terminal and C-terminal to it. The final output from this search was sorted to give those proteins that had a predicted even number of TMDs prior to the twin-arginine motif and that had a predicted single TMD immediately following the twin-arginine motif (available as supplementary online file at: http://www.lifesci.dundee.ac.uk/groups/tracy_palmer/docs/CombinedTATFindTMHMMoutput.docx). We subsequently manually searched this list to identify any proteins with a predicted C-terminal cofactor-binding domain.

From the output we identified a further actinobacterial Rieske homolog from *Kitasatospora setae* (KSE_30950) that is predicted to have five TMDs, with the twin arginine motif adjacent to TMD5. We also identified two further families of predicted metalloproteins that shared features of dual-inserted proteins (shown schematically in Fig 4A). Sco3746, also from S. *coelicolor* is predicted to have five TMDs, with a predicted molybdenum cofactor (MoCo) binding domain at the C-terminus and conserved histidine residues in TMDs 2, 3 and 4 that are predicted to co-ordinate two heme *b* moieties (Fig 4A). The twin arginine motif, which is conserved across homologous proteins (Fig S4) directly precedes TMD5. Homologues of Sco3746 were identified across the actinobacteria, as well as in firmicutes, chloroflexi and euryarchaeota, and each carries a twin arginine motif directly preceding TMD5 (Examples from each phyla are shown in Fig S4). Protein Q1NSB0 from the delta proteobacterium *MLMS-1* is also predicted to have five TMDs and to contain seven 4Fe-4S clusters, three at the cytoplasmic side and four at the extracellular side of the membrane (Fig 4A; Fig S5). Again the conserved twin arginine motif directly precedes TMD5 and homologues of this protein are encoded in many prokaryotic genomes including those from the chloroflexi, nitrospirae and euryarchaeota phyla (Fig S5).

**Figure 4:**
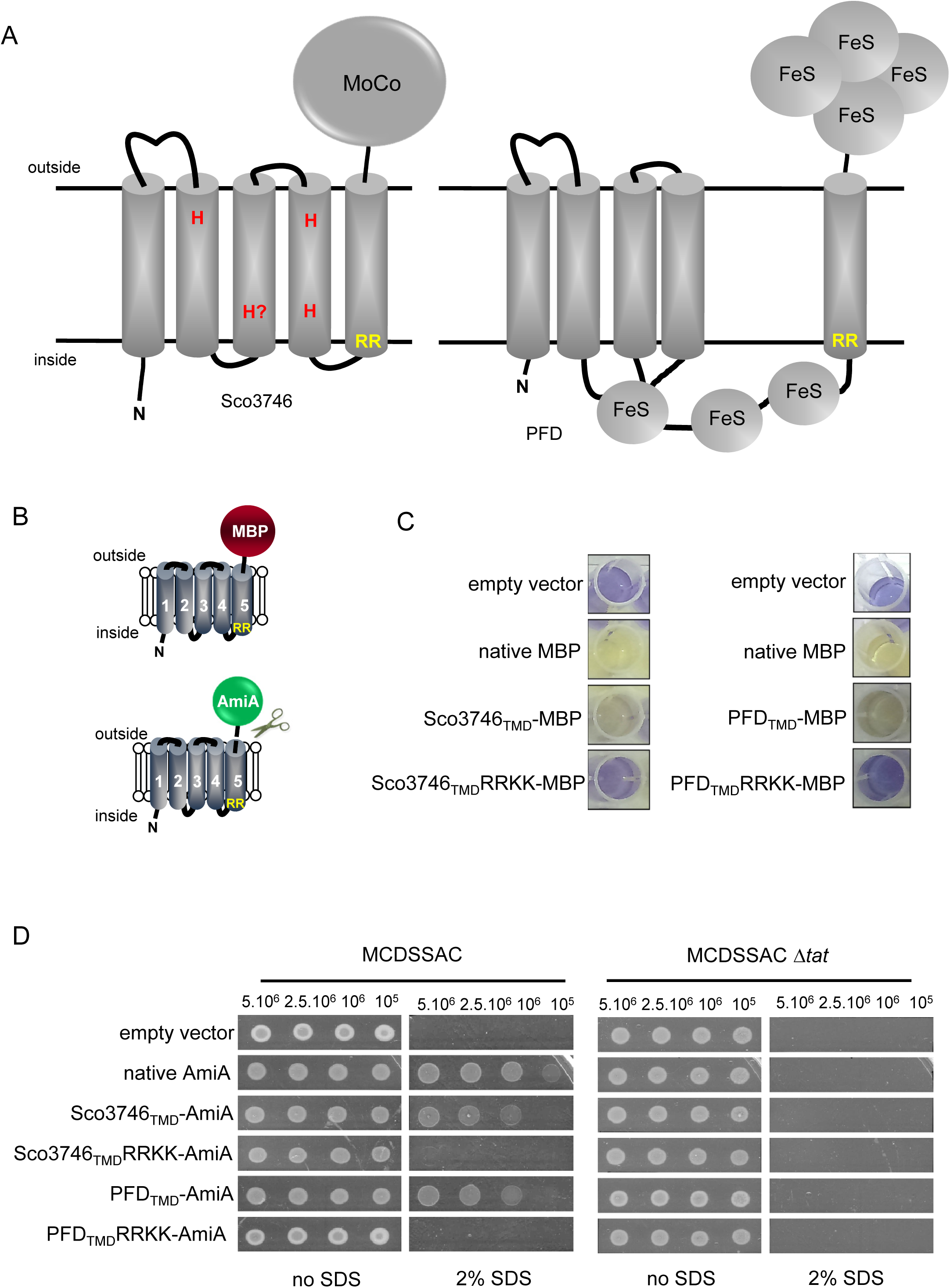
Two further families of Tat-dependent polytopic membrane proteins. (A) Schematic representation of a polytopic predicted molybdenum cofactor (MoCo) binding protein (Sco3746, left) and a polytopic polyferredoxin (PFD, right) identified bioinformatically as candidate dual-inserted membrane. The twin arginines of the Tat recognition sequence are highlighted in yellow. Four histidines in the TMDs of Sco3746 and homologues that are predicted to ligate two *b* hemes are shown in red. Note that three of these histidines are conserved (Fig S4) but the one at the N-terminal end of TMD3 (H?) is not. (B) Fusions of the TMDs of Sco3746 and of Q1NSB0 (PFD) to maltose binding protein (MBP) or AmiA are shown as cartoons. As before, a signal peptidase I cleavage site (indicated by scissors) was introduced at the end of predicted TMD5 to allow release of AmiA from the membrane (22). (C) *E. coli tat*^+^ strain HS3018-A (that lacks chromosomally-encoded MBP) harboring pSU18 (empty vector), or the same vector producing native MBP, Sco3746_TMD_-MBP, PFD_TMD_-MBP or the twin-arginine substituted variants Sco3746_TMD_RRKK-MBP and PFD_TMD_RRKK-MBP, as indicated, was cultured overnight, resuspended in minimal medium containing 1% Maltose and 0.002% Bromocresol purple. Cells were diluted either 2.5 fold (left hand panel) or 10 fold (right hand panel) in the same medium and incubated without shaking at 37°C for 24h (left hand panel) or 48h (right hand panel). (D) *E. coli* strains MCDSSAC or an isogenic *tatABC* mutant harboring pSU18 (empty vector), pSU18 producing native AmiA, or pSU18 producing either Sco3746_TMD_-AmiA or PFD_TMD_-AmiA fusion proteins, or variants of these where the twin-arginine motif was substituted to twin lysine (Sco3746_TMD_RRKK-AmiA/PFD_TM_DRRKK-AmiA) were serially diluted and spotted onto LB or LB containing 2% SDS. The plates were incubated for 20 hours at 37°C.

### Reporter proteins fused to Sco3746 or predicted polyferredoxin from *MLMS-1* are translocated by the Tat pathway

To confirm that these newly identified proteins were indeed Tat substrates, we designed constructs whereby the predicted five TMDs of Sco3746 or *MLMS-1* polyferredoxin (PFD; cloned as a synthetic gene) were fused to the reporter proteins AmiA or maltose binding protein (MBP; Fig 4B; exact positions of the fusions are shown in Figs S4 and S5). As shown in Fig 4C, *E. coli malE*^-^ cells harboring MBP fused to these regions of either protein decolorized maltose minimal medium containing the pH indicator dye bromocresol purple. This indicates that the MBP portion of the fusion protein has been translocated to the periplasmic side of the membrane. To confirm that this translocation was dependent on the Tat pathway, the twin-arginines of the Tat recognition motif were substituted for two lysines. This conservative substitution abolished maltose fermentation (Fig 4C), indicating that MBP translocation was dependent on the Tat pathway. Similar findings were made using the AmiA reporter fusions. Fig 4D shows that, as expected, when either plasmid-encoded Sco3746_TMD_-AmiA or PFD_TMD_-AmiA was produced in the *tat*^+^ strain lacking native AmiA/C, growth on SDS was supported. Export was dependent on the Tat pathway since growth on SDS was not supported in the *tat* strain, or in the *tat*^+^ strain if the twin arginine motif was substituted for twin lysine. We conclude that Sco3746 and PFD are dependent on the Tat pathway for their assembly.

### Sco3746_TMD_ and PFD_TMD_ fusions are stably inserted in the membrane in the absence of a functional Tat system

We next determined whether these fusion proteins were stably inserted into the membrane. Fig 5A shows that both Sco3746_TMD_-MBP and PFD_TMD_-MBP were detected exclusively in the in the membrane fraction of a *tat*^+^ strain at close to their theoretical masses (68 kDa for Sco3746TMD-MBP and 81 kDa for PFD_TMD_-MBP). It should be noted that the relatively poor expression of PFD_TMD_-MBP necessitated long exposure times for visualisation by western blot, thus two additional non-specific bands were also detected by the MBP antibody for these samples. Substitution of the Tat consensus arginine pair for di-lysine did not detectably affect the amount of fusion proteins produced, nor their membrane localization, indicating that membrane insertion of each of these fusions occurred independently of the Tat system. This was confirmed by repeating the analysis in a *tat*^-^ strain, where as expected the fusions were again detected exclusively in the membranes. Washing the membranes with 4M urea did not extract either protein (Fig 5B), indicating that they were integrally inserted into the membrane in the absence of the Tat pathway. This indicates the participation of a second protein translocase, almost certainly the Sec pathway, in the insertion of these proteins into the membrane.

**Figure 5:**
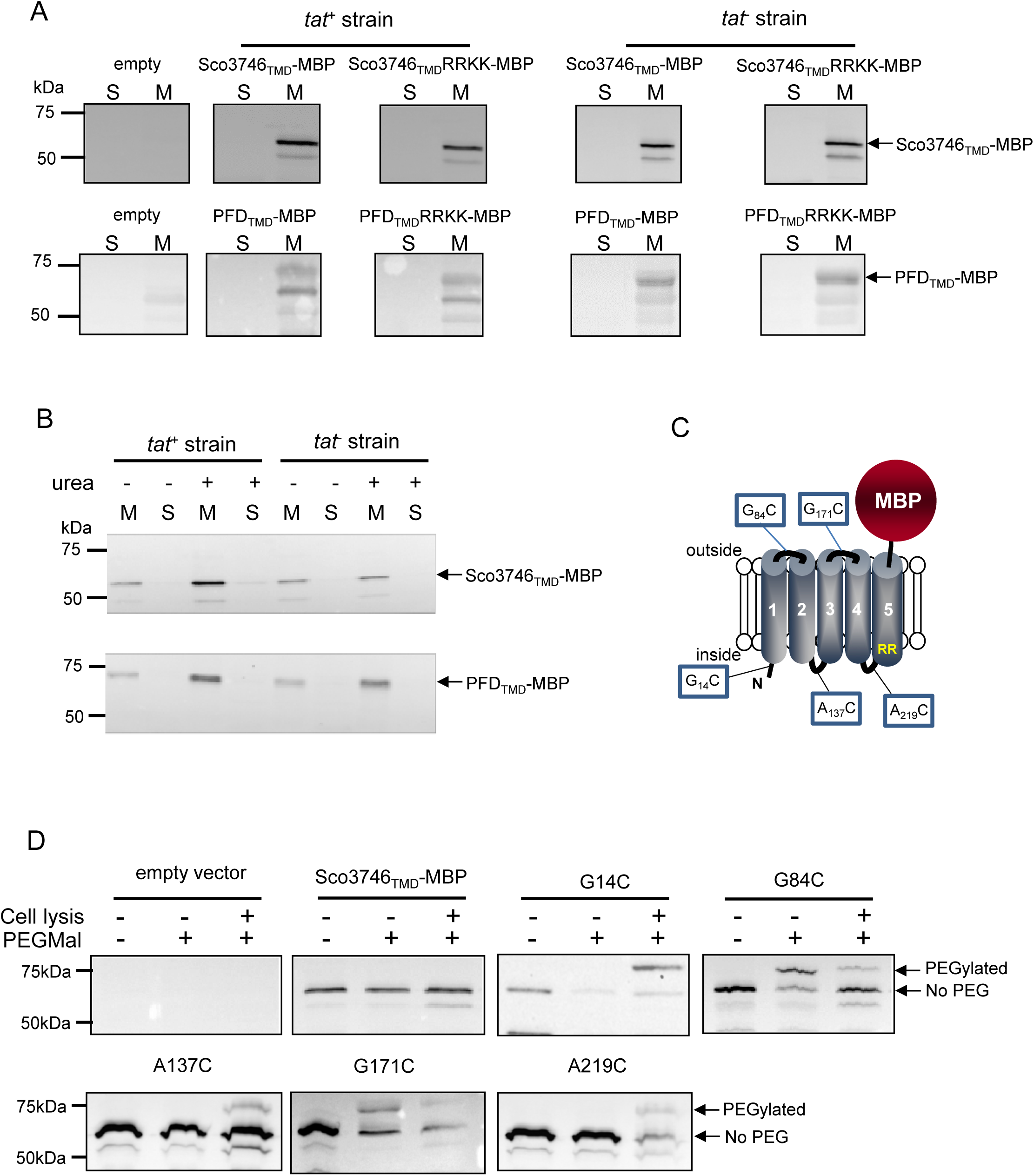
Topological analysis and membrane integration of CO3746_TMD_-MBP and PFD_TMD_-MBP. (A) Membrane (M; 100μg protein) and soluble (S; 50μg protein) fractions *of E. coli* HS3018-A (ΔmalE, *tat*^+^) and HS3018-AΔtat strains harboring pSU18 (empty vector), pSU18 encoding Sco3746_TMD_-MBP or PFD_TMD_-MBP fusion proteins, or variants of these where the twin-arginine motif was substituted to twin-lysine were separated by SDS-PAGE (12% acrylamide), transferred to nitrocellulose membrane and immunoblotted with an anti-MBP antibody. (B) Crude membranes of the same strains and plasmids were treated with 4M urea, and the presence of the fusion proteins in the wash supernatant (S) and pelleted membrane (M) was analyzed by immunoblotting as in (A). (C) Predicted locations of cysteine substitutions of Sco3746_TMD_-MBP used for topology analysis. (D) Cell suspensions of strain HS3018-A harboring pSU18 alone (empty vector), or pSU18 encoding Sco3746_TMD_-MBP or the indicated single cysteine substitutions of Sco3746_TMD_-MBP were incubated with buffer alone, with 5mM MAL-PEG, or were lysed by sonication and incubated with 5mM MAL-PEG. Subsequently all samples were quenched, lysed and membranes pelleted by ultracentrifugation. Membrane samples (150 μg of protein) were separated by SDS PAGE and immunblotted as in (A).

### Sco3746_TMD_-MBP has five TMDs

To confirm the predicted topology of the hydrophobic domain of Sco3746, we undertook a cysteine accessibility study. The Sco3746_**TMD**_-MBP fusion is naturally devoid of cysteine residues. Guided by topology prediction programs we made three Cys substitutions (G14C, A137C and A219C) that are predicted to reside at the cytoplasmic side of the membrane and two (G84C and G171C) that are located in predicted extracellular loops (Fig 5C). We produced these constructs in a *tat^+^* strain and probed cysteine accessibility using the reagent methoxypolyethyleneglycol maleimide (MAL-PEG). This reagent, which has a mass of around 5000 Da, can pass through the outer membrane in the presence of EDTA, but is impermeable to the inner membrane. Fig 5D shows that the G84C and G171C variants of Sco3746_TMD_-MBP clearly labelled with MAL-PEG in whole cells confirming that they are extracellular. By contrast, G14C, A137C and A219C variants were not labelled in whole cells but were labelled upon cell lysis, consistent with them having a cytoplasmic location. Taken together we conclude that the Sco3746_TMD_ portion of the Sco3746_TMD_-MBP fusion has 5 TMDs.

### A conserved mechanism regulates Sec-Tat transfer for three dual-targeted protein families

Our prior results analysing the interaction of actinobacterial Rieske proteins with the Sec pathway indicated that a combination of low hydrophobicity of the Tat-dependent TMD coupled with the presence of positive charges close to the C-terminal end of that TMD promoted release of the polypeptide from the Sec pathway. We therefore inspected the sequences of Sco3746 homologues and of PFD proteins to see whether these features are conserved across protein families. Fig S4 shows that several non-conserved positive charges are located close to the C-terminus of TMD5 of the Sco3746 homologues examined, and analysis of predicted ΔG_app_ values for membrane insertion of the five TMDs (Table 4) shows a positive ΔG_app_ for TMD5 suggesting that it may potentially be poorly recognised by Sec. Interestingly, unlike the actinobacterial Rieske proteins which have a highly conserved loop region between Sec-dependent TMD2 and Tat-dependent TMD3, the Sco3746 homologs have non-conserved loop sequences between TMD4 and TMD5 that show apparent length variability (although all of them are predicted to be at least 8aa long, which is the minimum loop length we defined for efficient recognition of Sco2149 TMD3 by the Tat pathway; Table 2).

**Table 4.** Predicted ΔG_app_ values (in kcal mol^-1^) for membrane insertion of each of the five TMDs of the indicated predicted MoCo-binding proteins analysed using the ΔG_app_ prediction server (http://dgpred.cbr.su.se/) and based on hydrophobicity scales generated from (36, 69). This server uses the SCAMPI2/TOPCONS servers (67, 68) to predict the positions of the TMDs and for *S. coelicolor* Q9L0V6 (Sco3746) predicts TMD1 to span aa 39-61, TMD2 to span aa 99-121, TMD3 to span aa 139-161, TMD 4 to span aa 173-194 and TMD5 to span aa 223-242.

| Family/Species | Protein ID | Predicted $\Delta G_{app}$ | | | | |
| --- | --- | --- | --- | --- | --- | --- |
|  |  | TM1 | TM2 | TM3 | TM4 | TM5* |
| *Phycococcus sp.* | WP_056916643.1 | -1.666 | -0.199 | -1.593 | 1.595 | 0.964 |
| *Solirubrobacter soli* | WP_053225688.1 | -1.778 | 1.394 | -2.239 | 1.681 | 1.135 |
| *Alicyclobacillus acidocaldarius* | C8WRE4 | -0.540 | -1.802 | -1.128 | 0.143 | 0.231 |
| *Ktedonobacter sp.* | OLB54713.1 | -0.479 | -2.647 | -2.375 | -1.257 | 1.762 |
| *Haloprofundus marisrubri* | WP_058581003.1 | 1.067 | -1.424 | -0.421 | -2.107 | 1.299 |
| *Streptomyces coelicolor* | Q9L0V6 | -2.268 | -0.011 | 0.328 | 0.512 | 1.376 |
| *S. coelicolor* G234L, S235L |  |  |  |  |  | -0.757 |
| *S. coelicolor* G234L, S235L, M239L, F2440L |  |  |  |  |  | -1.240 |

\*Bla is fused to *S. coelicolor* Sco3746 (Q9L0V6) after amino acid 247 after amino acid 252 (full sequence of all of the fusion proteins used in this study can be found in Supplementary Information).

We constructed ‘short’ (after aa 252) and ‘long’ (after aa 272) variants of Sco3746_TMD_ fused to Bla (Fig 6A), and expressed these in a *tat*^-^ strain to score for Sec-translocation of TMD5. Fig 6B shows that for the short fusion there is some degree of insertion of TMD5 by the Sec pathway because the M.I.C. for ampicillin mediated by this construct is significantly higher than the basal level. Substitution of hydrophobic leucines into TMD5 is predicted to shift the ΔG_app_ for membrane insertion of TMD5 from positive to negative (Table 4), and indeed, substitution of two or more leucine residues into the short fusion doubled the M.I.C. for ampicillin (Fig 6B), consistent with an increased level of insertion of TMD5 by Sec. The long Sco3746_TMD_-Bla fusion harbours an additional positive charge relative to the short fusion (Fig 6A). Fig 6B shows that this extension reduced the M.I.C. for ampicillin almost to the level of the empty vector control, consistent with the positive charges at the C-terminal end of Sco3746 TMD5 modulating interaction of this TMD with the Sec pathway.

**Figure 6:**
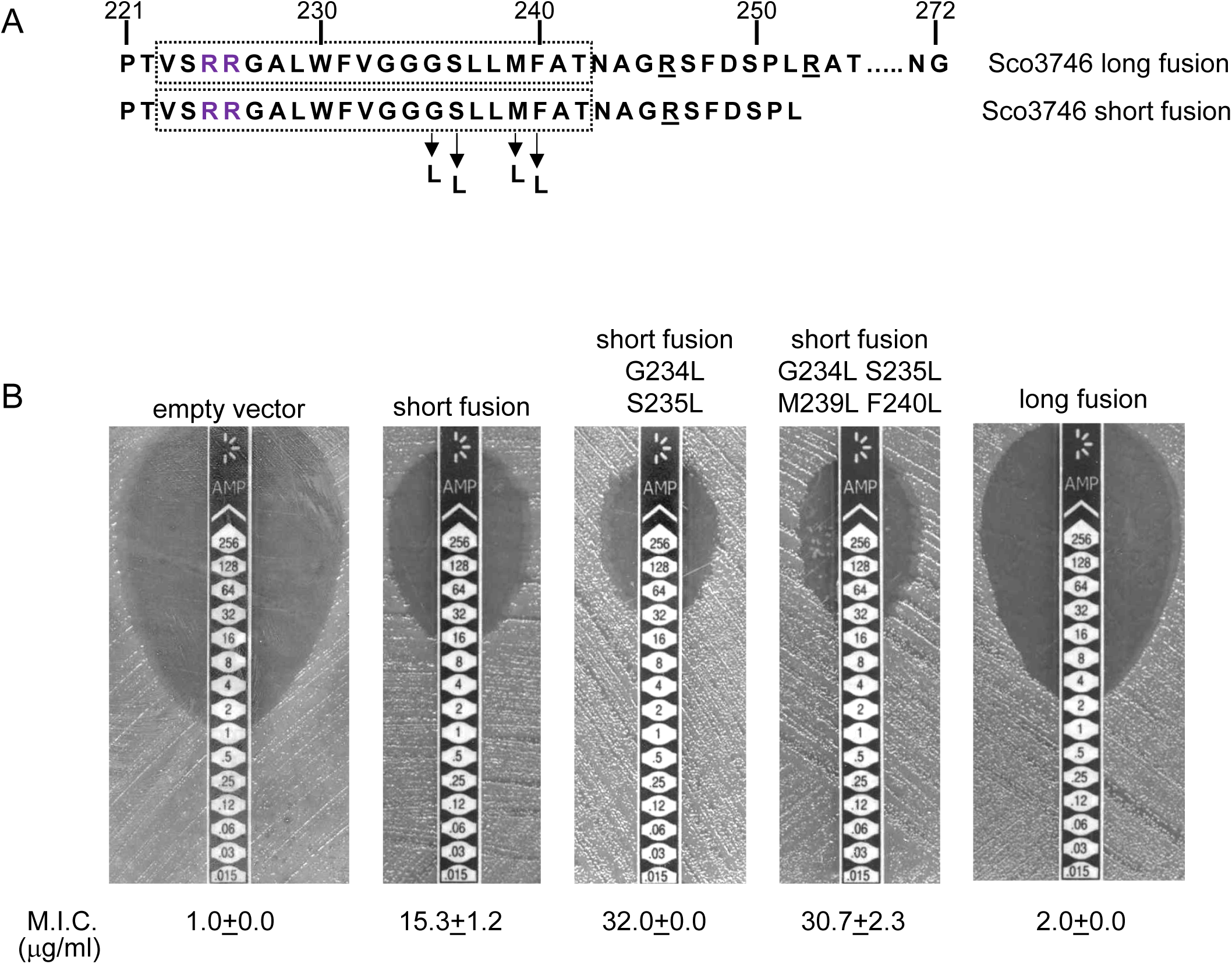
Relative hydrophobicity of TMD5 coupled with C-terminal positive charges modulate interaction of Sco3746 with the Sec pathway. (A) The sequence flanking TMD5 of Sco3746. The lower sequence extends to the position of the ‘short’ Sco3746-Bla fusion, and the amino acids in TMD5 substituted for leucine in this construct are shown. Top is the sequence fused to Bla in the ‘long fusion’. The predicted position of TMD5 was determined using the SCAMPI2/TOPCONS servers (67, 68) and is shown boxed. The twin arginines are shown in purple and positively charged amino acids C-terminal to TMD5 are underlined. (B) Representative images of M.I.C.Evaluator™ strip tests of strain DADE (*tat*^-^) harbouring pSU18 producing the indicated variants of Sco3746_TMD_-Bla In each panel the mean M.I.C ± s.d. is given at the bottom of each test strip (where *n*=3 biological replicates for each strain).

Similar to Sco3746 homologues, all of the PFD proteins analysed in Fig S5 also have a nonconserved positively charged region at the C-terminal end of TMD5. Analysis of predicted ΔG_app_ values for membrane insertion of the 5 TMDs was complicated by the observation that the iron-sulfur cluster binding regions were variably called as TMDs by some prediction programs. We therefore analysed only the first and fifth (Tat-dependent) TMDs (Table 5), and again it can be seen that the Tat-dependent TMD has a positive predicted ΔG_app_.

**Table 5.** Predicted ΔG_app_ values (in kcal mol^-1^) for membrane insertion of the first and last TMDs of the indicated predicted polyferredoxin proteins analysed using the ΔG_app_ prediction server (http://dgpred.cbr.su.se/) and based on hydrophobicity scales generated from (36, 69). This server uses the SCAMPI2/TOPCONS servers (67, 68) to predict the positions of the TMDs and for delta proteobacterium *MLMS-1* Q1NSB0 (PFD) predicts TMD1 to span aa 9-31 and TMD5 to span aa 338-359.

| Family/Species | Protein ID | Predicted $\Delta G_{app}$ | |
| --- | --- | --- | --- |
|  |  | TM1 | TM5* |
| *Latescibacteria*<br>*bacterium DG_63* | A0A0S7XC05 | -1.792 | 0.371 |
| *Nitrospirae bacterium*<br>*GWA2_46_11* | OGW21092.1 | -1.062 | 1.222 |
| *Caldithrix* sp.<br>*RBG_13_44_9* | OGB67066.1 | -3.560 | 1.438 |
| *Thermoplasmatales*<br>*archaeon DG-70-1* | KYK31751.1 | -1.474 | 1.715 |
| *Anaerolinea*<br>*thermolimosa* | WP_062193756.1 | -2.317 | 1.107 |
| *Anaeroarcus*<br>*burkinensis* | WP_027938385.1 | -0.975 | 0.836 |
| delta proteobacterium<br>*MLMS-1* | Q1NSB0 | -2.382 | 0.297 |
| delta proteobacterium<br>*MLMS-1* G354L,<br>R358L |  |  | -0.701 |
| delta proteobacterium<br>*MLMS-1* G354L,<br>P355L, R358L |  |  | -1.741 |

\*Bla is fused to delta proteobacterium *MLMS-1* PFD (Q1NSB0) after amino acid 364 (full sequence of all of the fusion proteins used in this study can be found in Supplementary Information).

To probe PFD interaction with the Sec pathway we designed ‘short’ (fused after aa 371) and ‘long’ (fused after aa 374) fusions of PFD_TMD_ from delta proteobacterium *MLMS-1* to Bla (Fig 7A) and produced these in a *tat*^-^ strain to score for Sec-translocation of TMD5. However, neither of these constructs mediated detectable export of β-lactamase as the M.I.C. for ampicillin was almost indistinguishable from the negative control (Fig 7B,C). We attribute this to the relatively poor expression of the PFD fusion proteins (e.g. Fig 5A). Next we substituted two, or three, leucine residues into TMD5 of PFD in each of the fusions, which is predicted to lower the ΔG_app_ value for TMD5 membrane insertion (Table 5). In agreement with this, Fig 7B shows that these substitutions significantly increased the level of interaction of the short fusion with the Sec pathway, giving mean M.I.C.s for ampicillin of 9.3μg/ml for the G354L, R358L and 12.0μg/ml for the G354L, P355L, R358L substitutions, respectively. These same substitutions also increased the interaction of the long fusion with Sec as they also conferred some resistance to ampicillin, each giving a mean M.I.C. of 6.7 μg/ml (Fig 7C). However it is clear that the same leucine substitutions confer lower levels of resistance to ampicillin when they are present in the long construct than when they are in the short construct (compare Fig 7B with Fig 7C). Since the shorter construct harbours one less positive charge at the C-terminal end of TMD5 we conclude that the additional positive charge present in the extended fusion reduces the level of membrane insertion by Sec. Taken together our results demonstrate that the mechanism of Sec release of the final TMD is conserved across all known families of dual Sec-Tat targeted membrane proteins.

**Figure 7:**
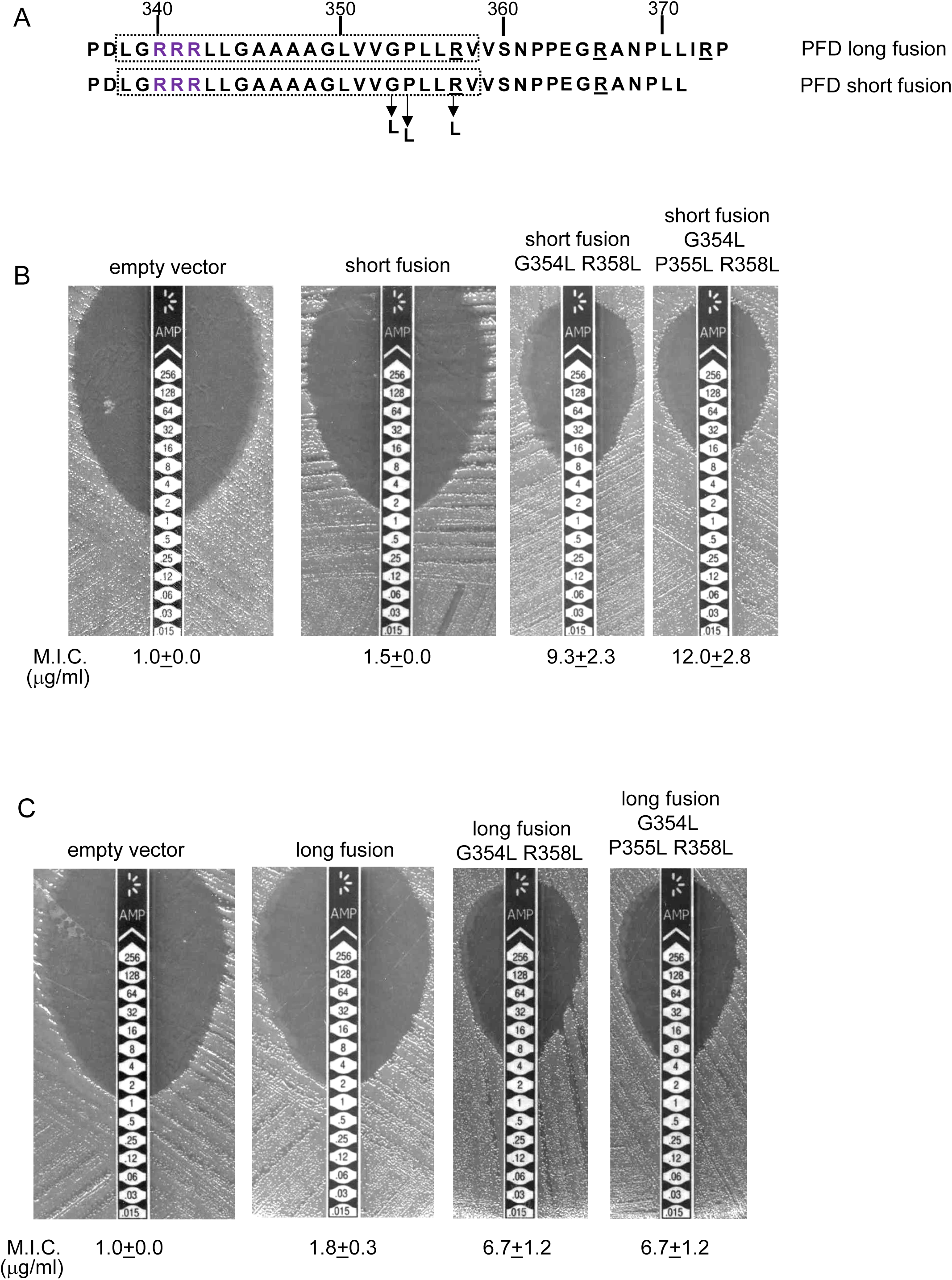
Relative hydrophobicity of *MLMS-1* PFD TMD5 coupled with C-terminal positive charges governs interaction with the Sec pathway. (A) The sequence flanking TMD5 of *MLMS-1* PFD. The lower sequence (corresponding to ‘short fusion’ in parts B and C) extends to the position of the shorter PFD-Bla fusion, whereas the top sequence is the sequence fused to Bla in the ‘long fusion’. The predicted position of TMD5 was determined using the SCAMPI2/TOPCONS servers (67, 68) and is shown boxed. The twin arginines are shown in purple and positively charged amino acids C-terminal to TMD5 are underlined. Amino acids in TMD5 substituted for leucine in both constructs are shown. (B and C) Representative images of M.I.C.Evaluator™ strip tests of strain DADE (*tat*^-^) harbouring pSU18 producing the indicated variants of PFD_TMD_-Bla In each panel the mean M.I.C ± s.d. is given at the bottom of each test strip (where *n*=3 biological replicates for each strain).

## Discussion

In a previous study we identified the actinobacterial Rieske FeS protein as the first protein known to be targeted to the plasma membrane by the dual action of the Sec and Tat translocases. The mechanism by which translocation is coordinated between the two pathways was not known, although a length- and sequence-conserved loop region between Sec-dependent TMD2 and Tat-dependent TMD3 was implicated in this process (22). Intensive investigation into the principles governing the correct biogenesis and topology of membrane proteins has revealed that the relative hydrophobicity of a TMD along with the location of positively charged amino acids are key features that govern the insertion and orientation of transmembrane segments (34, 36, 40). Here we show that that these principles are exploited by nature to regulate translocation of Rieske by the Sec pathway and allow its hand-off to Tat prior to insertion of the final TMD. None of the features of the highly conserved loop region, other than the presence of one or more positively charged amino acids that serve as topology signals, plays any discernible role in co-ordinating the Sec and Tat pathways and may therefore be required for cofactor insertion or interaction with other components of the cytochrome *bc*_1_complex.

A bioinformatic analysis of prokaryotic genome sequences identified two further families of polytopic membrane proteins that share the predicted features of Sec-Tat dual-targeting. Each of these has five TMDs, with the fifth TMD immediately preceded by a consensus Tat recognition motif. A representative member of each family was shown to be membrane inserted through the action of two translocases, with the Tat system recognising the final TMD. Importantly, the low hydrophobicity of the final TMD coupled with C-terminal positive charges, identified through our analysis of the *S. coelicolor* Rieske protein as being critical for Sec-release, are conserved across these further protein families, and were confirmed experimentally to govern release of this final TMD from Sec. Thus a common mechanism is at play to orchestrate the integration of dual Sec-Tat targeted membrane proteins.

A model for how such proteins are assembled is shown in Fig 8, using the actinobacterial Rieske protein as an example. According to the model, the Sec-dependent helices are inserted co-translationally. The positively-charged twin-arginines N-terminal to the final TMD imposes an N-in, C-out orientation on this helix. However, the C-terminal positive charges prevent the full insertion of this TMD because the relatively low hydrophobicity is insufficient to drive translocation of the C-terminal positively charged region (45, 46). This is experimentally supported by our findings that substitution of a single leucine residue into TMD3 of the *S. coelicolor* Rieske-Bla fusion is sufficient to greatly increase its Sec-dependent insertion despite the presence of two positive charges at the C-terminal end. Accordingly, it is likely that this final TMD is released by the Sec pathway as a re-entrant loop. Following folding of the cofactor-containing domain, the Tat machinery mediates translocation of the folded domain across the membrane, releasing the Tat-dependent TMD into the bilayer.

**Figure 8:**
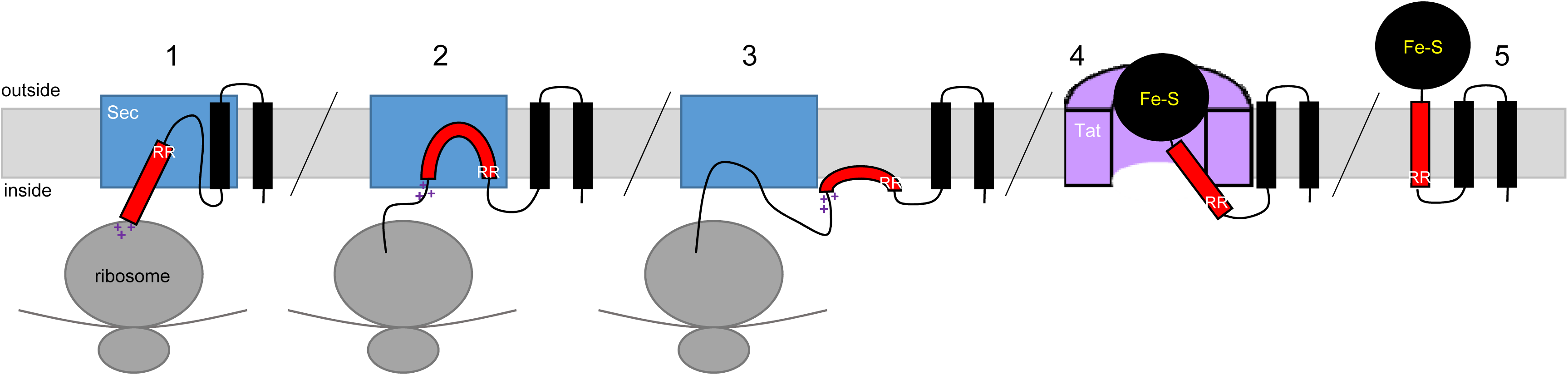
Model for actinobacterial Rieske protein assembly. 1. TMDs 1 and 2 are inserted into the membrane cotranslationally by the Sec machinery (blue box). The Sec machinery interacts with TMD3 in an N-in, C-out orientation. 2. The positive charges at the C-terminal end of TMD3 force an orientational preference on the helix and it is not inserted by the Sec machinery. 3. The hydrophobic segment of TMD3 is released from the Sec machinery as a re-entrant loop. As there are no further TMDs within the Riekse sequence the Sec machinery releases the polypeptide. 4. Once translation is complete the iron-sulfur cluster is inserted into the protein and the Tat machinery (pink half-cylinder) interacts with TMD3 to translocate the folded globular domain across the membrane. 5. The fully assembled Rieske protein is released into the membrane to interact with partner proteins.

Signal peptides of soluble Tat substrates often contain one or more positively-charged residues in their c-regions which are known to act as Sec-avoidance motifs. Removal of these charges results in signal sequences that can mediate efficient transport by the Sec machinery (41-43). Furthermore, signal peptides that direct proteins to the Tat machinery are known to be less hydrophobic than Sec signal peptides and if the hydrophobicity of a Tat signal peptide is increased it can also mediate efficient transport by the Sec pathway (42). However, despite possessing these ‘Sec-avoidance’ features, over half of the native *E. coli* Tat signal peptides are capable of transporting reporter proteins through the Sec pathway if fused to an appropriate passenger domain (47). This raises the possibility that rather than being an exception, Sec interaction with Tat signal peptides is much more frequent, and that following abortive attempts at Sec-translocation, membrane-associated twin-arginine signal peptides are common substrates of the Tat pathway. In this context it should be noted that both thylakoid and *E. coli* Tat substrates interact with the membrane before subsequent interaction with Tat machinery (48-51).

Our work has shown that dual targeted Sec-Tat dependent membrane proteins are dispersed across two domains including Gram-negative and Gram-positive bacteria and euryarchaea, indicating that the biogenesis of dual-targeted membrane proteins is a common feature of prokaryotes. It is interesting to note that distant homologs of both the predicted heme-Moco binding protein, Sco3746, and the *MLMS-1* polyferredoxin are widely found as separate polypeptides. For example *E. coli* MsrP/MsrQ (formerly YedY/YedZ encoded by *yedYZ*) are, respectively, a Sec-dependent polytopic heme *b* protein and Tat-targeted soluble MoCo-containing periplasmic protein that together use electrons from the respiratory chain to catalyse the repair of proteins containing methionine sulfoxide (52, 53). Likewise *MLMS-1* polyferredoxin is a fusion of NapH, a Sec-dependent polytopic protein with four TMD that coordinates [4Fe-4S] iron-sulfur clusters at the cytoplasmic side of the membrane, with NapG, a Tat-dependent periplasmic protein that is predicted to co-ordinate four [4Fe-4S] at the periplasmic side of the membrane. Collectively NapGH form a quinol dehydrogenase complex that in *E. coli* and *Wolinella succinogenes* is involved in nitrate respiration (54,55). The close relationship of such proteins and their corresponding genes raises the possibility that dual-targeted proteins arose during the course of evolution from separate polypeptides but adjacent genes. Alternatively, the ancestral proteins may have been single, dual-targeted polypeptides that subsequently separated in some organisms.

## Materials and methods

### Bacterial strains, Plasmid construction and growth conditions

All strains used in this study are derived from *Escherichia coli* K-12 and are listed in Table S1. Strain DH5α (Stratagene) was used for molecular biology applications. Strains MC4100 (56) and DADE (as MC4100; Δ*tatABCD,* Δ*tatE;* (57)) were used for work with Bla fusions, MCDSSAC (26) and MCDSSACΔtat (as MCDSSAC; Δ*tatABC*::Apra; (22)) were used for work with AmiA fusions, and HS3018-A (58) and HS3018-AΔtat (As HS3018-A; Δ*tatABCD,* Δ*tatE;* (22)) were used for work with MBP fusions.

The amino acid sequences of all of the fusion proteins used in this study can be found in Supplementary Information. All plasmids used and generated in this study are listed in Table S2 and all oligonucleotides are listed in Table S3. To generate pSU-PROM AmiA, DNA encoding full length AmiA was PCR amplified using oligonucleotides BamHI AmiA and SU18.2 with pSU18 AmiA (22) as a template, digested with *Bam*HI and *Hind*III and inserted into similarly digested pSU-PROM (28). To generate pSU-PROM Sco2149_TMD_-AmiA, the Sco2149_TMD_-AmiA allele was excised from pSU-TM123-AmiA (22) by digestion with *Bam*HI/*Hind*III and ligated into similarly digested pSU-PROM. To generate pSU-PROM Sco2149_TMD_-Bla, the *amiA* coding region was excised from pSU-PROM Sco2149_TMD_-AmiA by digestion with *Xba*I*/Hind*III and replaced with the coding sequence for the mature region of Bla obtained by PCR amplification from pBR322 that had been similarly digested. To extend the Sco2149_TMD_-Bla fusion to aa205 of Sco2149, the region covering Sco2149 codons 1-205 were amplified using oligonucleotides Sco2149_TMD_ and Sco2149_TMD_ extension and cloned as a *Bam*HI-*Xba*I fragment into similarly digested pSU-PROM Sco2149_TMD_-Bla to generate pSU-PROM Sco2149TMDextended-Bla.

DNA encoding the first 247 amino acids of Sco3746 was PCR amplified using oligonucleotides Sco3746For and Sco3746Rev with *Streptomyces coelicolor* M145 chromosomal DNA as a template, digested with *Bg/*II and *Xba*I and inserted into pSU-PROM (28) that had been digested with BamHI and XbaI. The region covering the *tat* promoter and Sco3746_TMD_ coding region was excised using *Eco*RI*/XbaI* and ligated into similarly digested pSU18 (59).

Subsequently DNA encoding the mature regions of AmiA (from pSU-PROM Sco2149_TMD_-AmiA) or MBP (from pTM123-MBP, (22) were cloned in as *Xba*I-*Hind*III fragments to give Sco3746_TMD_-AmiA and Sco3746_TMD_-MBP, respectively. To construct Sco3746_TMD_-Bla the first 252 amino acids of Sco3746 was PCR amplified using oligonucleotides Sco3746For and Sco3746(252)Rev with *S. coelicolor* M145 chromosomal DNA as a template, digested with *Bg/*II and XbaI and inserted into similarly digested pSU-PROM Sco2149_TMD_-Bla (thus replacing the Sco2149 coding sequence with Sco3746). Subsequently the *Xba*I site was replaced with *Kpn*I by Quickchange site-directed mutagenesis using oligonucleotides Sco3746_TMD_BlaFor and Sco3746_TMD_BlaRev. To extend the Sco3746_TMD_-Bla fusion to aa272 of Sco3746, the region covering Sco3746 codons 1-272 were amplified using oligonucleotides SU18.1 and Sco3746_TMD_extension and pSU18PROM Sco3746_TMD_-Bla as template. This was digested with *Eco*RI and *Kpn*I fragment and ligated into a similarly digested pSU18PROM Sco3746_TMD_-Bla as template to generate pSU18PROM Sco3746TMDextended-Bla.

A synthetic gene encoding the transmembrane region (residues 1-227) of the Rieske protein (QcrA) from *Mycobacterium tuberculosis* strain RV2195 was codon optimised for *E. coli* K12 expression (OPTIMIZER, (60)) and the synthetic gene was purchased ready cloned in pUC57 (GenScript). The MtbRieske_TMD_ coding region was subcloned by digestion *Rca*I-*Xba*I and ligated into pBAD24 (61) using vector sites *Nco*I/*Xba*I. It was then digested *Bam*HI/*Xba*I and ligated into pSU-PROM Sco2149_TMD_-Bla in place of Sco2149_TMD_. To extend the MtbRieske_TMD_- Bla fusion to aa243 of QcrA, the region covering QcrA 1-243 were amplified using oligonucleotides MtbRieske_TMD_ and MtbRieske_TMD_ extension and pBAD24-QcrA as template, digested with *Bam*HI-*Xba*I and ligated into similarly digested pSU-PROM MtbRieske_TMD_-Bla to generate pSU-PROM MtbRieske_TMD_extended-Bla.

The transmembrane coding region (residues 1-364) of the predicted polyferredoxin (PFD) from delta proteobacterium *MLMS-1* (NCBI GI:494503356) was codon optimised for *E. coli* K12 expression (OPTIMIZER, (60)) and the synthetic gene was purchased already cloned into pBluescript (Biomatik). The PFD_TMD_ coding region was excised with *RcaI/XbaI* and cloned into pBAD24 (61) that had been digested with *Nco*I/*Xba*I. Subsequently DNA encoding the mature region of MBP (excised from pTM123-MBP (22)) was cloned in as an *Xba*l-*Hind*III fragment. The entire PFD_TMD_-MBP coding region was subsequently excised as an *Eco*RI-*Hind*III fragment and cloned into similarly digested pSU18 (59) to give pSU18 PFD_TMD_-MBP. To construct pSU18 PFD_TMD_-AmiA, the MBP coding region was excised and replaced with the AmiA coding region (as an *Xba*I/*Hind*III fragment from pSU-PROM Sco2149_TMD_-AmiA). To construct the PFD_TMD_-Bla fusion (which covers up to aa371 of PFD), oligonucleotides SU18.1 and PFD_TMD_BlaRev were used to amplify the PFD coding sequence (with pSU18 PFD_TMD_-Bla as template). The product was digested with *Eco*RI and *Kpn*1 and ligated into similarly digested pSU18PROM Sco3746_TMD_-Bla to generate PFD_TMD_-Bla. The PFD_TMD_ coding sequence in this construct was further extended to residue 374 using oligonucleotides SU18.1 and PFD_TMD_ extension and pSU18 PFD_TMD_-Bla as template. The resultant product was digested with *Eco*RI-*Kpn*I and ligated into similarly digested pSU18 PFD_TMD_-Bla to generate pSU18 PFD_TMD_ extended-Bla.

Site-directed mutagenesis was performed using the QuickChange™ method (Stratagene) according to manufacturer’s instructions. Deletion mutants were generated from a modified QuickChange™ method adapted from (62). Briefly, forward and reverse primers were designed to remove up to 5 residues at a time, overlapping by 12 nucleotides upstream and downstream of the region to be deleted with an overhang of 12 nucleotides at either end. For truncations larger than 5 residues the template used, already contained a downstream deletion of all residues but the additional 5 residues to be removed. All constructs were verified by DNA sequencing.

Unless otherwise stated, *E. coli* strains were grown aerobically overnight at 37°C in Luria-Bertani (LB) broth supplemented with appropriate antibiotic/s at the indicated final concentrations - ampicillin (125 μg/ml), kanamycin (50 μg/ml), apramycin (25 μg/ml) and chloramphenicol (25 μg/ml). Filter-sterilised SDS solution was added to the media to final concentration of 1 to 2% as indicated. Phenotypic growth tests in the presence of SDS were performed as follows: overnight cultures were diluted to OD_600_ 0.1 and 5 μl aliquots were spotted in a serial dilution series from 10^4^ cells to 10^1^ cells per 5 μl for Sco2149_TMD_-AmiA and 5.10^6^ to 10^5^ for Sco3746_TMD_-AmiA and PFD_TMD_-AmiA on LB agar supplemented with 1 or 2% SDS. Phenotypic testing for maltose fermentation employed the approach of (22) using maltose-bromocresol purple broth prepared with M9 minimal medium supplemented with 0.002% bromocresol purple (Roth) and 1% maltose. Growth was performed in 96-well plates incubated without shaking for 24 h to 48 h at 37°C. *E. coli* susceptibility to ampicillin was determined by assessing the Minimum Inhibitory Concentration (M.I.C.) that prevented growth. Stationary phase cultures were diluted to OD_600_ 0.1 and LB agar plates were inoculated by swabbing the diluted culture to generate a lawn of bacteria. Oxoid M.I.C.Evaluator™ test strips (Thermo Fisher Scientific) containing a gradient of 0-256 μg/ml ampicillin were placed onto the lawn and incubated at 37°C for 18h. The M.I.C. value (in μg/ml) was read from the scale where the pointed end of the ellipse intersects the strip according to manufacturer’s instructions.

Photographs of 96-well plates were captured as JPG files using a digital camera (DX AF-S NIKKOR 18-55 mm; Nikon) and colonies on agar with a digital scanner (EPSON perfection 3490 PHOTO). JPG files were imported into Gimp for cropping but otherwise were not processed.

### Subcellular fractionation

Membrane and cellular fractions were prepared as described by Keller *et al.* (22) with modifications. *E. coli* cells were grown overnight at 37°C in LB medium with appropriate antibiotics, subcultured and harvested at OD_600_ of 0.2 for cells producing Sco2149 derivatives or OD_600_ of 0.5 for cells producing Sco3746 and PFD derivatives. Cells producing Sco2149 constructs were resuspended in the same volume of hypertonic buffer (20mM Tris-HCl pH7.5/200mM NaCl) supplemented with EDTA-free protease inhibitor (Roche). Cells producing Sco3746 or PFD constructs were diluted to give a final OD_600_ of 0.2 in the same buffer. Cells were then lysed by sonication (Branson Digital Sonifier 250) and the suspension was centrifuged for 10 min at 20 000 *g* at 4°C to remove unbroken cells and large cellular debris. The resulting supernatant was then ultracentrifuged for 1 hour at 220 000 *g* at 4°C to separate membrane and soluble fractions. An aliquot of the soluble fraction was kept for analysis and the membrane pellet was resuspended in 50mM Tris-HCl pH 7.5; 5mM MgCl_2_; 10% (v/v) Glycerol. Protein concentration was estimated by the Lowry method (63) using the DCTM Protein Assay kit (Bio-Rad) and a standard curve generated with Bovine Serum Albumin (BSA). Membrane and soluble fractions were snap-frozen and kept at -20°C until further analysis. Urea extraction was undertaken as described previously (22).

### Cysteine accessibility experiments

Cysteine accessibility was assessed as described by (64) with the following modifications. Cultures were harvested at OD_600_ of 0.8 and resuspended to give a final OD_600_ of 0.3 in a labeling buffer (50mM HEPES, 5mM MgCl_2_ pH 6.8) supplemented with EDTA-free protease inhibitor (Roche). An aliquot of the sample was lysed by sonication. Labeling was then performed for 1 h at room temperature on intact or lysed cells using 5mM methoxypolyethylene glycol maleimide (MAL-PEG) at room temperature for 1 h in the presence of 5mM EDTA. A control without addition of labeling reagent was systematically included. The reaction was quenched by addition of 100 mM dithiothreitol (DTT) and the whole cell samples were then lysed by sonication. All samples were subsequently centrifuged for 10 min at 20 000 *g*, 4°C to remove non-broken cells and the resulting supernatant ultracentrifuged for 30 min at 200 000 *g* at 4°C to pellet the membranes. The membrane pellet was resuspended in 60 μl of 50mM Tris-HCl pH 7.5; 5mM MgCl_2_; 10% (v/v) glycerol for analysis by SDS PAGE and western blotting.

### Protein analysis

Proteins were separated by Tris-glycine SDS-PAGE (7.5%, 10%, 12% or 15% polyacrylamide, as indicated) and transferred onto nitrocellulose membrane either with semi-dry (TransBlot SD SemiDry Transfer Cell, Bio-Rad) or dry transfer (iBlot2, Life technologies). Proteins were detected with primary antibodies raised against either Sco2149 (a monoclonal Sco2149 peptide antibody generated by GenScript against the N-Terminal epitope CLPPHEPRVQDVDER), Bla (monoclonal antibody, Abcam ab12251), MBP (monoclonal antibody, NEB E8032L), BamA (polyclonal antibody, (65)). Bands were revealed with chemiluminescence (Clarity™ Western ECL Blotting Substrate, Biorad) after incubation with secondary antibody coupled to HRP (anti-mouse IgG or Anti-Rabbit IgG, Biorad). Light-emitting bands were visualised with a CCCD camera (GeneGNOME XRQ Syngene). ImageJ (66) was used for densitometry analysis. The density of Sco2149-associated signals were normalised against the BamA-associated signals which was used as a loading control. The results were expressed as percentage of the normalised signal obtained for the unmodified Sco2149_TMD_-AmiA or Sco2149_TMD_-Bla fusion proteins.

### Bioinformatic analysis

A Perl script was used to run all proteins from all completed prokaryotic genomes available in Genbank at the time of analysis through the TATFind program (version 1.4; (44)) and the TMHMM program (version 2.0c; (2)). For each protein, the outputs of two were combined to arrive at the position of twin arginine motif, and the number of transmembrane helices on the N-terminal side, and the C-terminal side, respectively, of the twin arginine motif. Only those proteins which had an even number of predicted TMDs N-terminal to the twin arginine motif and only one TM helix C-terminal to it, were kept and can be found in supplementary information.

## Acknowledgements

This work was supported by the UK Biotechnology and Biological Sciences Research Council (through grants BB/L000768/1 and BB/J004561/1), and the UK Medical Research Council (through grant G0901653). TP is a Wellcome Trust Investigator. We thank Rebecca Keller, Jeanine de Keyzer, Arnold Driessen, Gunnar von Heijne and Frank Sargent for helpful discussion and advice, Hannah Garnham for her assistance with constructing some of the cysteine substitutions used in this study and Felicity Alcock for critical reading of the manuscript.

## Author Contributions

F.J.T., M.B., G.C., G.B. and T.P. designed research; F.J.T., M.B., G.C. and G.B. performed research; F.J.T., M.B., G.C., G.B. and T.P. analyzed data; and F.J.T., M.B. and T.P. wrote the paper.

